# In Silico Design and Analysis of Genetic Circuit-Based Whole-Cell Biosensors for Detecting Groundwater Contaminants

**DOI:** 10.1101/2020.12.22.424054

**Authors:** Samuel Fajemilua, Solomon Bada, M. Ahsanul Islam

**Affiliations:** Department of Chemical Engineering, Loughborough University, Loughborough, Leicestershire LE11 3TU, UK

**Keywords:** Whole-cell biosensor, genetic circuit, genetic design automation, iBioSim, groundwater contaminants

## Abstract

Contaminants of emerging concern (CEC) such as tetracycline, erythromycin, and salicylic acid in groundwater can seriously endanger the environment and human health due to their widespread and everlasting harmful effects. Thus, continuous monitoring of various CEC concentrations in groundwater is essential to ensure the safety, security, and biodiversity of natural habitats. CECs can be detected using whole-cell biosensors for environmental surveillance and monitoring purposes, as they provide a cheaper and more robust alternative to traditional and expensive analytical techniques. In this study, various genetic circuit designs are considered to model three biosensors using the genetic design automation (GDA) software, iBioSim. The genetic circuits were designed to detect multiple CECs, including atrazine, salicylic acid, and tetracycline simultaneously to produce quantitative fluorescent outputs. The biosensor responses and the viability of the genetic circuit designs were further analysed using ODE-based mathematical simulations in iBioSim. The designed circuits and subsequent biosensor modelling presented here, thus, not only show the usefulness and importance of GDA tools, but also highlight their limitations and shortcomings that need to overcome in the future; thereby, providing a practical guidance for further improvement of such tools, so that they can be more effectively and routinely used in synthetic biology research.

## Introduction

The subsurface groundwater aquifers are enormous storehouses of water and one of the Erath’s most valuable resources. These aquifers provide groundwater in usable quantities for world’s vast majority of people, as their uses range from agricultural use such as irrigation for crop production to industrial and public water supply use. Around 25% of the UK’s public water supply comes from groundwater [1]. In North America alone, around 150 million people rely on groundwater for their drinking water supply [2]. Similarly, 85% of drinking water supplies are provided by groundwater in developing countries such as India [3]. On account of this, interest in the groundwater environment has heightened, and resources are being administered to ensure and maintain its safety. Of these resources, many are focused on the removal of contaminants which mainly originate from agricultural, industrial, and municipal activities [4], [5]. Some of these contaminants have been reported to cause a range of health problems, including cancer [6], diabetes [7], and birth defects [8]. Accordingly, the detection and removal of these contaminants are of great importance for the safety and security of not only these valuable groundwater aquifers but also of the environment and public health in general.

Over the past decade, certain chemical compounds that were previously present at insignificant levels in groundwater have become increasingly detectable. These chemical compounds have been referred to as contaminants of emerging concerns (CECs) [9]. CECs are commonly found in pesticides, herbicides, fungicides, and pharmaceutical compounds [10]. The pharmaceutical CECs, such as tetracycline, erythromycin, and salicylic acid have been shown to increase the rates of causing different types of cancers, organ damage, and antibiotic resistance [11]–[13]. Similarly, the CECs commonly found in pesticides, including atrazine, allethrin, and carbaryl have been shown to induce biodiversity damage, pest resistance, and endocrine disruption [13]–[16]. In addition, Eriksen *et al* [17] observed a 30% 40% increase in prostate cancer due to the CEC, perfluorooctanoate. Further, trihalomethanes present in groundwater were associated with an increased risk of spontaneous abortions [18]. However, due to the limited number of studies and contradictions in study outcomes, the connection between CECs and health risks have not yet been fully established. Further research is required to fully elucidate this connection.

Previously, groundwater contaminants were detected by means of geophysical methods such as NMR geotomography [19] and spectroscopic techniques such as UV lasers [20]. In the past ten years, significant advances in synthetic biology research have been achieved [21]. These advances have resulted in the use of biosensors to detect contaminants such as arsenic [22], toluene [23], and xylene [24]. Biosensors utilise biological responses and physiochemical components to generate a measurable signal for detection of an object [25]. Depending on the transducing mechanism used, these devices can exist in many forms such as electrochemical biosensors, optical biosensors, and resonant biosensors [26]. However, the development cost and operating boundaries associated with these biosensors have prompted questions on their feasibility for environmental surveillance and monitoring applications. The advances in synthetic biology [27], together with the cost pressures and operating boundaries, have provoked notable interests in the development of whole-cell biosensors due to their low cost and versatility [28].

Whole-cell biosensors incorporate microorganisms to detect specific pollutants by receiving signals and processing these signals via genetic circuits to produce a quantitative or qualitative output. Genetic circuits can process signals by exploiting transcriptional regulation to orchestrate the gene expression [28]. Another technique is through the incorporation of Boolean logic gates in the cell [29]. Boolean logic gates make use of the logic functions such as AND, OR, NOT, and all of their combinations [30]. Once the input requirements of these logic functions are met in the gates, an output is executed. Whole-cell biosensors tend to detect a wider range of substances when compared to conventional biosensors [31]. Moreover, whole-cell biosensors can operate in much harsher pH values and temperatures than its conventional counterparts [31]. Although whole-cell biosensors boast great advantages, synthetic biologists may face challenges when designing feasible biosensors, as issues may arise when implementing these designs in the field. For this reason, the whole-cell biosensors have not yet reached their potential. Accordingly, attention has been directed towards the optimisation and practicality of whole-cell biosensor designs [32].

Preceding efforts to construct whole-cell biosensors to detect groundwater contaminants are similar to that of Liu *et al* [33]. A number of studies have further demonstrated the scalability of whole-cell biosensors, detecting a range of contaminants [22]–[24]. However, the studies focused on single-input biosensors rather than considering the multi-input options. Multi-input biosensors are capable of handling complex environments such as groundwater aquifers, and this can be achieved through the integration of Boolean logic gates [34]. Moreover, biosensors for the detection of CECs have been poorly studied in comparison to other groundwater contaminants.

Recent advances in computational synthetic biology has allowed the development of computational models of genetic circuits to be implemented in a whole-cell biosensor. Over the past decade, various software tools have been developed to aid the advancement of synthetic biology designs. One such software is the genetic design automation (GDA) tool, iBioSim [35], [36]. iBioSim provides an environment whereby visualisation, modelling, and analysis of genetic circuits can be achieved; thus, permitting the analysis of various circuit design options in terms of robustness, operating conditions, and operating boundaries [35]. This study uses iBioSim to (1) design three genetic circuits capable of detecting three CECs: atrazine, salicylic acid, and tetracycline, both individually and simultaneously to produce measurable florescent outputs, (2) model and simulate these genetic circuits in the context of biological cells to emulate how the whole-cell biosensors work, and (3) analyse the results obtained from the modelling and simulation of the designed genetic circuits as biosensors. Furthermore, the factors to be considered for experimental implementation of the biosensor models will be discussed, and the drawbacks which may arise will be underscored.

## Materials and methods

### Genetic Circuit Design and Construction

To increase consistency and replicability, all genetic circuit designs proposed in this study were designed, modelled, and simulated in iBioSim [37]. An example of the circuit design procedure has also been described in detail in the supplementary information.

A genetic circuit usually consists of several biological parts, including promoters, ribosome binding sites (RBS), proteins, and terminators. A promoter is the specific DNA region which initiates the transcription of a gene [38]. Translation is then initiated at the RBS whereby messenger RNA (mRNA) is decoded to produce a protein [39]. The terminator is a section of DNA which stops the transcription process [40]. All biological parts were obtained from SynBioHub [41] and were designed following the BioBricks™ standards [42]. Figure 1 illustrates a genetic circuit designed for the detection of salicylic acid using standard parts from the Registry of Standard Biological Parts (for example, BBa_B0015 is the BioBricks name of a double terminator) and expressed in Synthetic Biology Open Language Visual (SBOL Visual) format. SBOL is an open-source standardised data format for sharing and exchanging in silico genetic circuit designs [43]. SBOL Visual was developed as an extension of SBOL to promote coherency when expressing the function and structure of genetic designs [44]. In Figure 1, the coding sequence codes for the red fluorescent protein, RFP (BBa_I766101), where coding for this protein is activated by the salicylic acid-sensitive promoter, P_Sal1_ (BBa_J61051). The ribosome binding site used to mediate the protein synthesis is RBS30 (BBa_K1789002), and the double terminator (BBa_B0015) was employed to end gene transcription. The various standard biological parts used from the parts registry to design the biosensors are summarised in Table 1. The detail SBOL Visual design for all biosensors can be found in the supplementary information.

**Figure 1.**
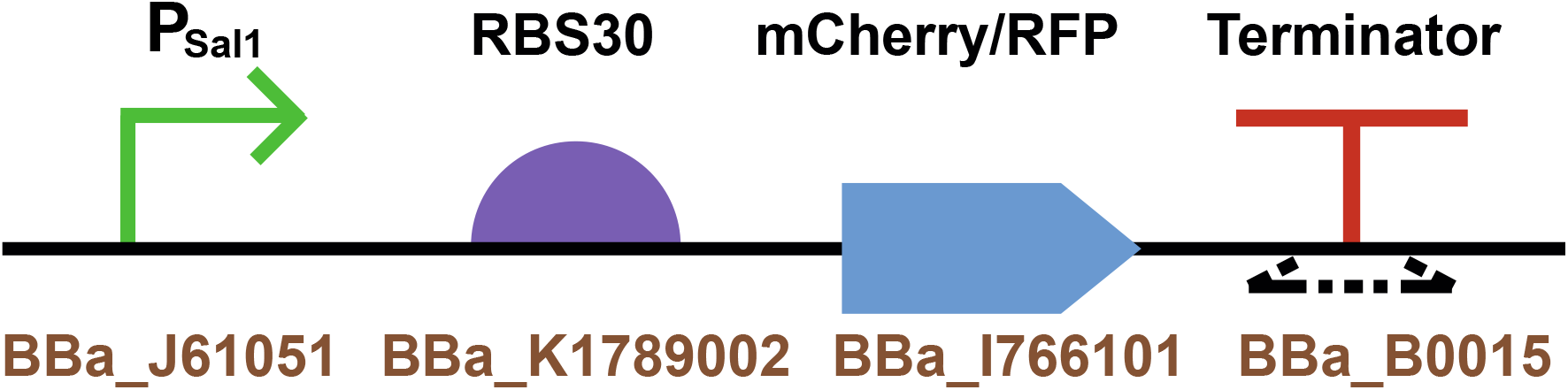
SBOL Visual design of a genetic circuit to detect salicylic acid. P_Sal1_ (BBa_J61051) is the promoter, RBS30 (BBa_K1789002) is the ribosome binding site, mCherry/RFP (BBa_I766101) is the reporter protein, and BBa_B0015 is the terminator.

**Table 1:**
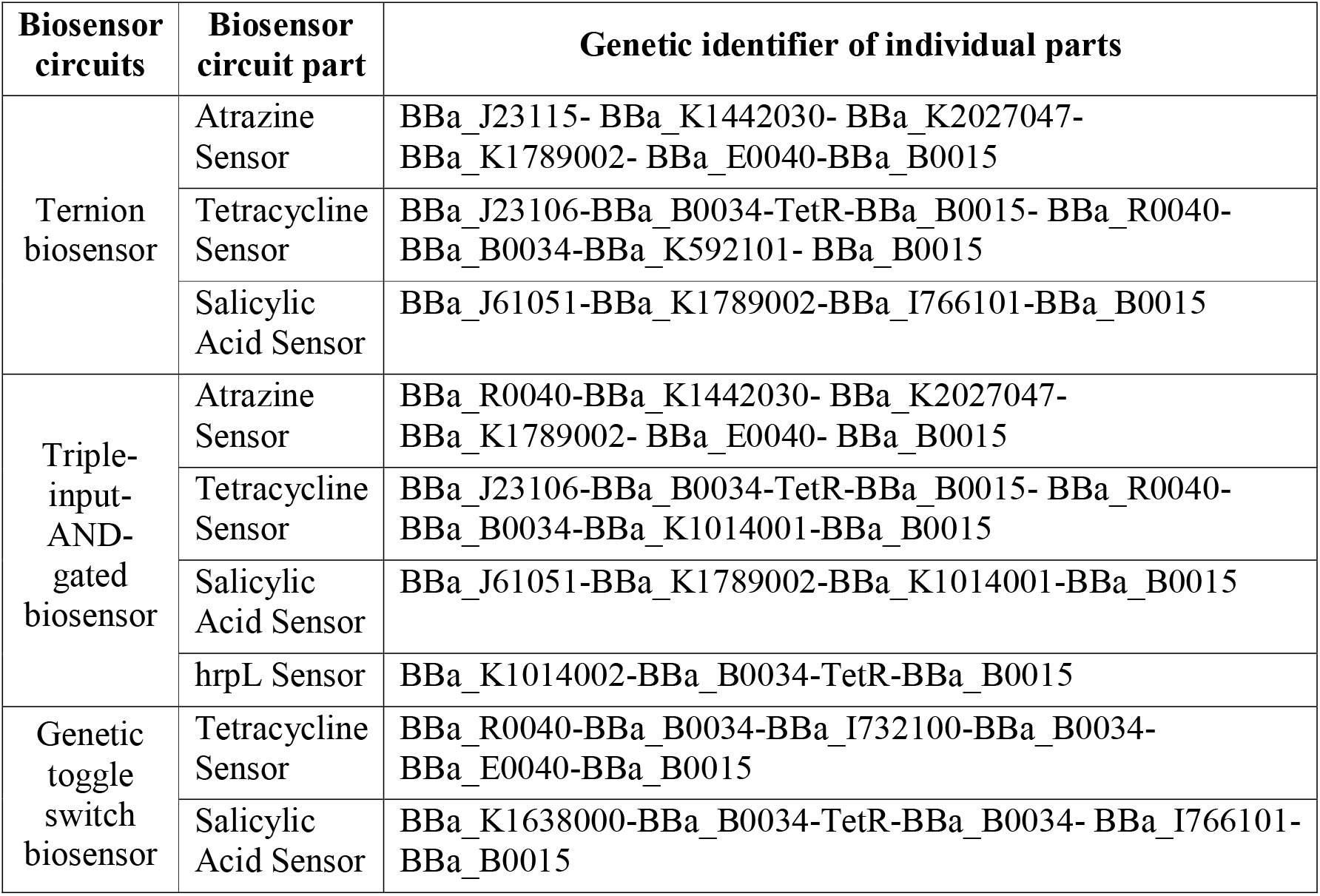
Summary of standard biological parts used to design different biosensor circuits

### Modelling biosensor circuits in iBioSim

The genetic circuit designs for simulating three different types of whole-cell biosensors in iBioSim are shown in Figure 2 (Fig. 2A is the ternion biosensor, Fig. 2B is the triple-input-AND-gated biosensor, and Fig. 2C is the genetic toggle switch biosensor). The ternion biosensor (Fig. 2A) is comprised of a three-cell consortium containing one sub-circuit for each cell. The sub-circuits are placed in Cell_1, Cell_2, and Cell_3 to detect atrazine, tetracycline, and salicylic acid for producing green fluorescent protein (GFP), yellow fluorescent protein (YFP), and RFP signals, respectively. The model also includes eight species (blue ovals in Fig. 2A) and four complex formation reactions (light green squares in Fig. 2A); species represent chemical components such as compounds, genes, and proteins, while complex formation reactions represent the chemical reactions transforming these species. These reactions are further described by four sub-models showing the formation of C1 TetR_production, C2 TetR_YFP, C3 Atrazine_GFP, and C4 Salicyclic_acid_RFP (Fig. 2A). Further details of all sub-models are shown in the supplementary information. Next, events (dark green rectangles in Fig. 2), representing biosensor responses to varying concentration of atrazine, tetracycline, and salicylic acid, were added to the models. In the case of Cell_1, the atrazine concentration was changed through the events, Atrazine_High and Atrazine_Low to simulate high and low atrazine concentrations in the environment.

**Figure 2.**
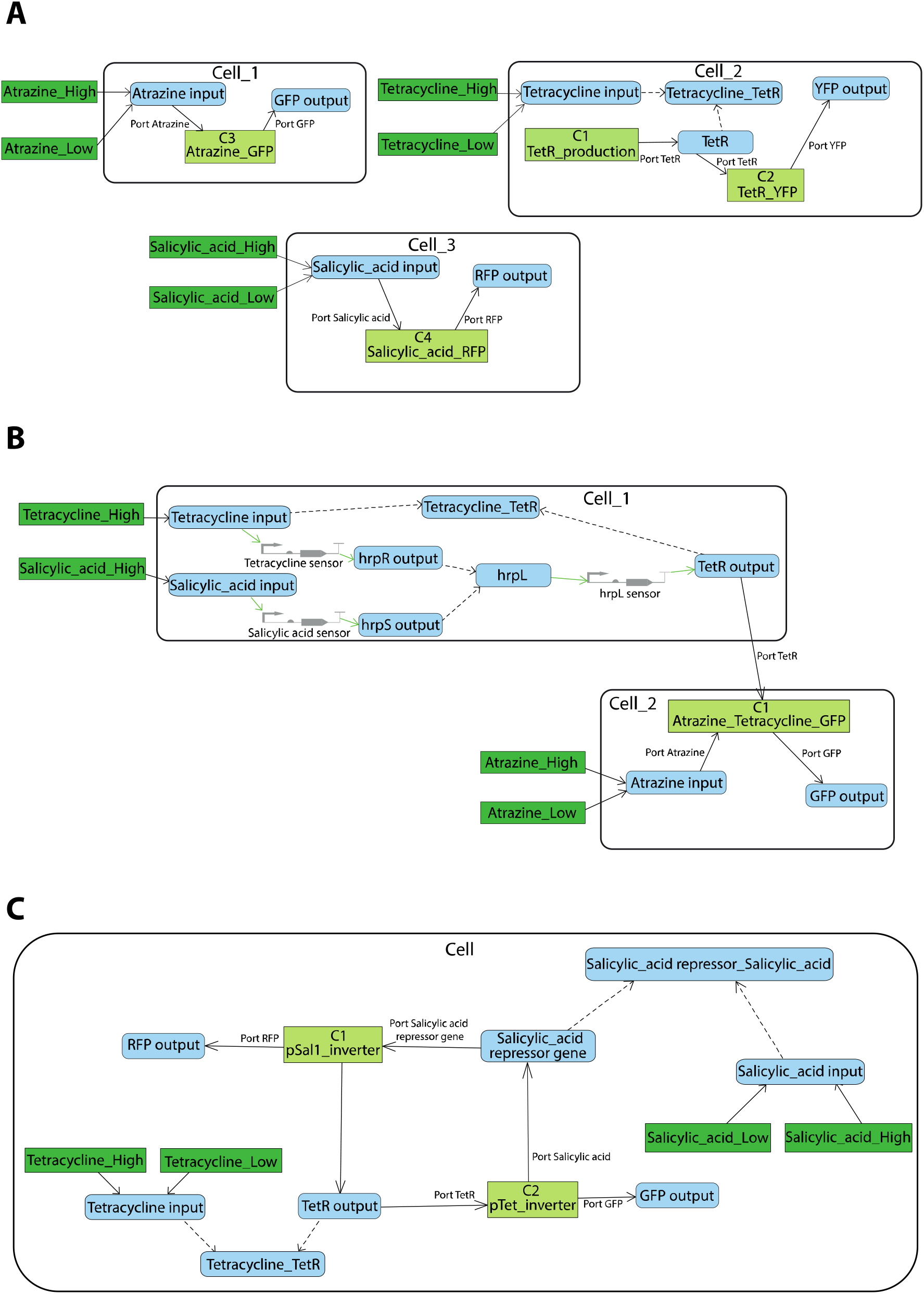
iBioSim models of the designed genetic circuits. (A) iBioSim model of the ternion biosensor circuit. The model includes eight species, four sub-models, six events, and two complex formation reactions. (B) iBioSim model of the triple-input-AND-gated biosensor circuit. This model has nine species, one sub-model, four complex formation reactions, four events, and three grey SBOL parts. (C) iBioSim model of the toggle switch biosensor circuit. This model includes eight species, two sub-models, four complex formation reactions, and four events. Species, sub-models, complex formation reactions, and events are represented by blue ovals, light green squares, dark green rectangles, and dashed arrows, respectively.

Similarly, tetracycline and salicyclic acid concentration changes were simulated in Cell_2 and Cell_3, respectively (Fig. 2A). The triple-input-AND-gated (Fig. 2B) and the genetic toggle switch (Fig. 2C) biosensor models were also simulated in iBioSim following the same approach as described for the ternion biosensor model.

### Mathematical basis of the designed biosensor circuits

Ordinary differential equations (ODEs) were used to simulate the biosensors in iBioSim. The Runge-Kutta-Fehlberg method was used to find numerical approximations to solve the ODEs [35]. To mathematically model the biosensors, the underlying reactions associated with gene expression must be deduced. The reactions in relation to the conversion of DNA to mRNA to protein can be written as [45]:

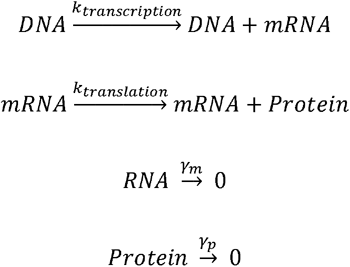

Where *k*_*transcription*_ and *k*_*translation*_ represent the rates of conversion of DNA to mRNA and mRNA to protein, respectively, while *γ*_*m*_ and *γ*_*p*_ are the degradation rates of mRNA and protein, respectively.

Thus, the rate of change of mRNA and protein level over time can be expressed using the following ODEs [45]:

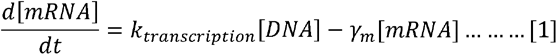

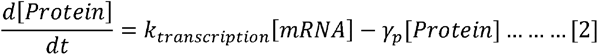

Before using the above differential equations for simulating biosensor responses, it is essential to formulate a mathematically accurate depiction of how a transcription factor regulates the gene expression. When modelling genetic circuits, the approach commonly taken emanates from the mass action kinetics [45]. According to this principle, the rate of a reaction in the genetic circuits is expressed as the product of a rate constant◻multiplied by the concentration of the inducer [46].◻In order to use this kinetic scheme for a genetic circuit, the following assumptions are required [45], [47]:

1. Polymerase binding, ribosome binding, and other intricacies of molecular interactions are ignored to reduce the complexity of the model,
2. The transcriptional activity is considered at steady state to accommodate for rapid change of transcriptional activity by virtue of the input signals, and ◻
3. The spatial effects are neglected to avoid partial differential equations. ◻

These assumptions led to the final form of the equations used to model the synthetically constructed biosensors. In the example of the genetic toggle switch, this would be:

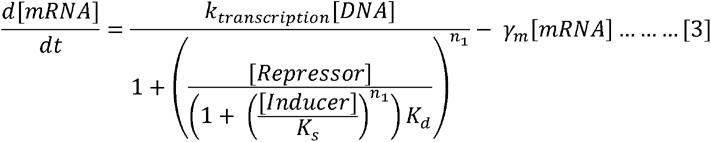

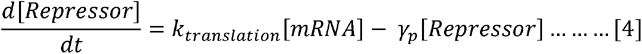

Where 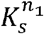 and 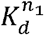 are constants. This permits the development of a system of ODEs for the genetic toggle switch leading to the following final solution:

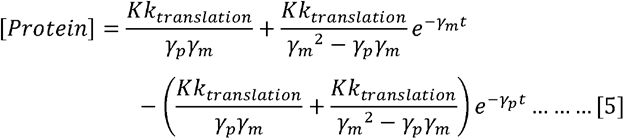

Where *K* can be defined as:

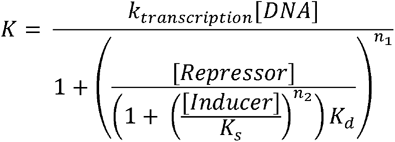

The full derivation of the genetic toggle switch is shown in the supplementary information.

## Results and Discussion

### Design and construction of biosensors

#### 1. Ternion biosensor

A ternion whole-cell biosensor was modelled using a three-cell consortium to detect atrazine, salicylic acid, and tetracycline by producing green, red, and yellow fluorescent signals as outputs, respectively (Figure 3). While the natural signalling mechanisms of host cells were used to detect salicylic acid and tetracycline, a synthesised riboswitch was used to provoke the transcription and translation of GFP in the presence of atrazine. The schematics of different sub-circuits of the ternion biosensor are shown in Figures 3A, 3B, and 3C. For the atrazine genetic circuit (Fig. 3A), the constitutive promoter, BBa_J23115 is used to continuously activate the atrazine riboswitch. Initially, the RBS is blocked to prevent the translation of GFP. In the presence of atrazine, an aptamer-atrazine bond is created, which unblocks the RBS to allow the transcription and translation of GFP to produce green fluorescence signals (Fig. 3A). In contrast to the atrazine genetic circuit, the salicylic acid sensor utilises a salicylic acid-sensitive promoter, P_Sal1_ (Fig 3B). In the presence of salicylic acid, the promoter activates the transcription and translation of RFP, producing red fluorescent signals. This incident demonstrates the detection of salicylic acid by the production of red signals as the output. The tetracycline genetic circuit (Fig. 3C) shows a constitutive promoter, BBa_J23106, continuously expressing the tetracycline-sensitive repressor, TetR. In the absence of tetracycline, TetR represses the tetracycline-sensitive promoter, P_Tet_; thereby, inhibiting the production of YFP. However, the presence of tetracycline results in a tetracycline-TetR complex formation leading to the reduction of TetR-P_Tet_ binding affinity. Consequently, P_Tet_ activates the transcription and translation of YFP to produce yellow fluorescent signals as the output; thus, achieving the detection of tetracycline by the ternion biosensor.

**Figure 3.**
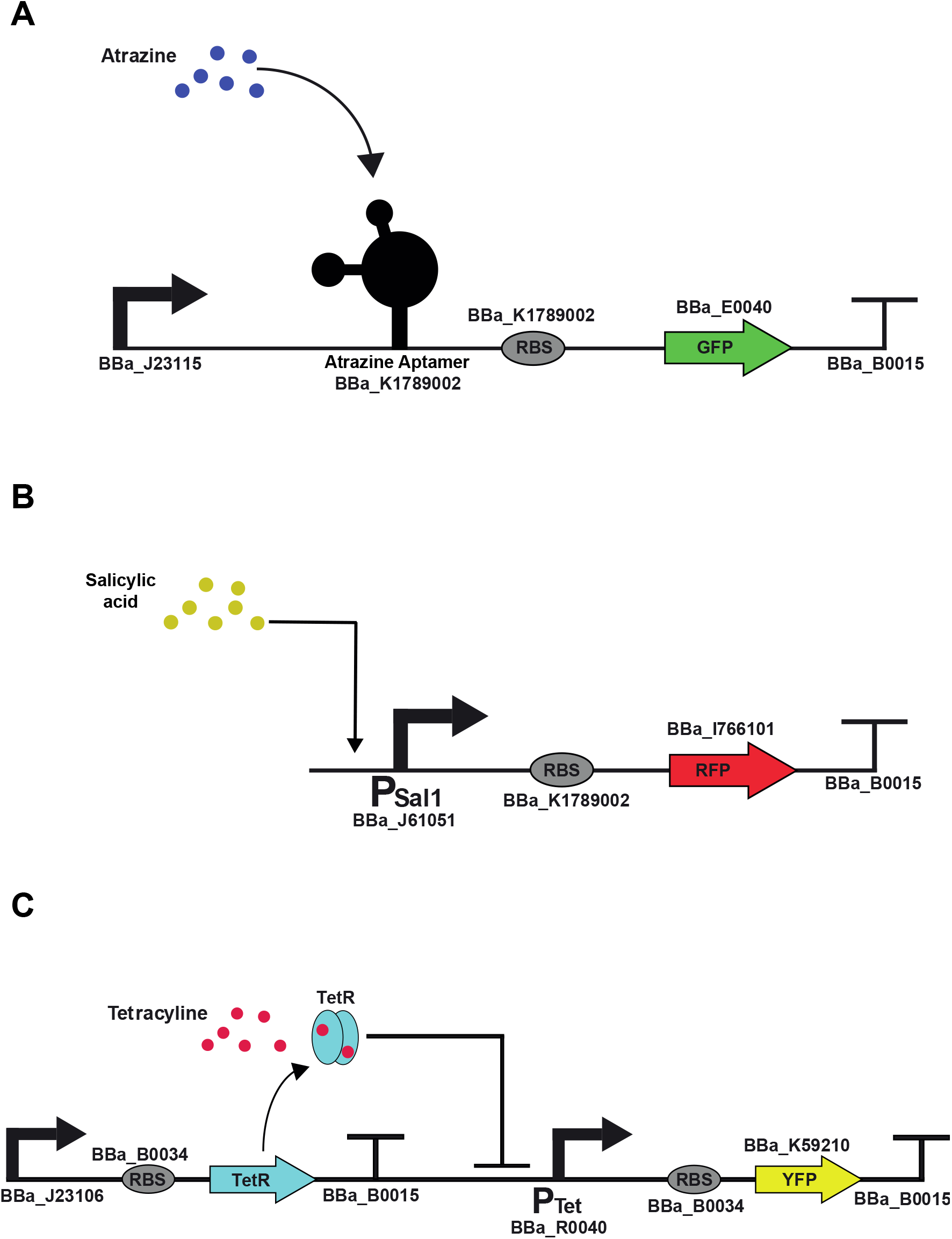
Schematic of the genetic circuits for the ternion biosensor model. The model contains the atrazine, salicylic acid, and tetracycline sensors shown in (A), (B), and (C) to generate green, red, and yellow signals by producing green fluorescent protein (GFP), red fluorescent protein (RFP), and yellow fluorescent protein (YFP), respectively.

#### 2. Triple-input-AND-gated biosensor

In situations such as groundwater contamination detection, demand for accuracy and specificity is prominent. Sensor systems with the ability to implement complex functions are essential to detect various contaminants in parallel [29]. Taking this into account, a biosensor making use of the Boolean logic function, AND was designed and modelled. For the triple-input-AND-gated biosensor (Figure 4), a two-cell consortium containing three genetic circuits was utilised to produce a single green fluorescent output. The first cell includes two genetic circuits, sensing the detection of tetracycline and salicylic acid, while the second cell contains only the atrazine detecting circuit. The sensors are of a similar structure to that of the ternion biosensor, the only difference being their respective output signals. For this biosensor model, the outputs for the detection of tetracycline and salicylic acid are the production of hrpR and hrpS proteins, respectively. Both hrpR and hrpS are enhancer-binding proteins that act as positive regulatory factors for P_hrpL_, the promoter for the alternative sigma factor, hrpL; thus, having the capability to activate it [48]. Upon activation of P_hrpL_, the TetR protein is produced by the first cell. TetR has the ability to diffuse across cells to form a complex with tetracycline molecules [49]. The tetracycline-TetR complex is capable of activating the P_Tet_ promoter in the second cell. Thus, in the presence of atrazine, both P_Tet_ and the atrazine riboswitch are activated to produce GFP, generating green fluorescent signals as the output (Figure 4). Therefore, the generation of green signals by the triple-input-AND-gated biosensor indicates the presence of all three contaminants in the system.

**Figure 4.**
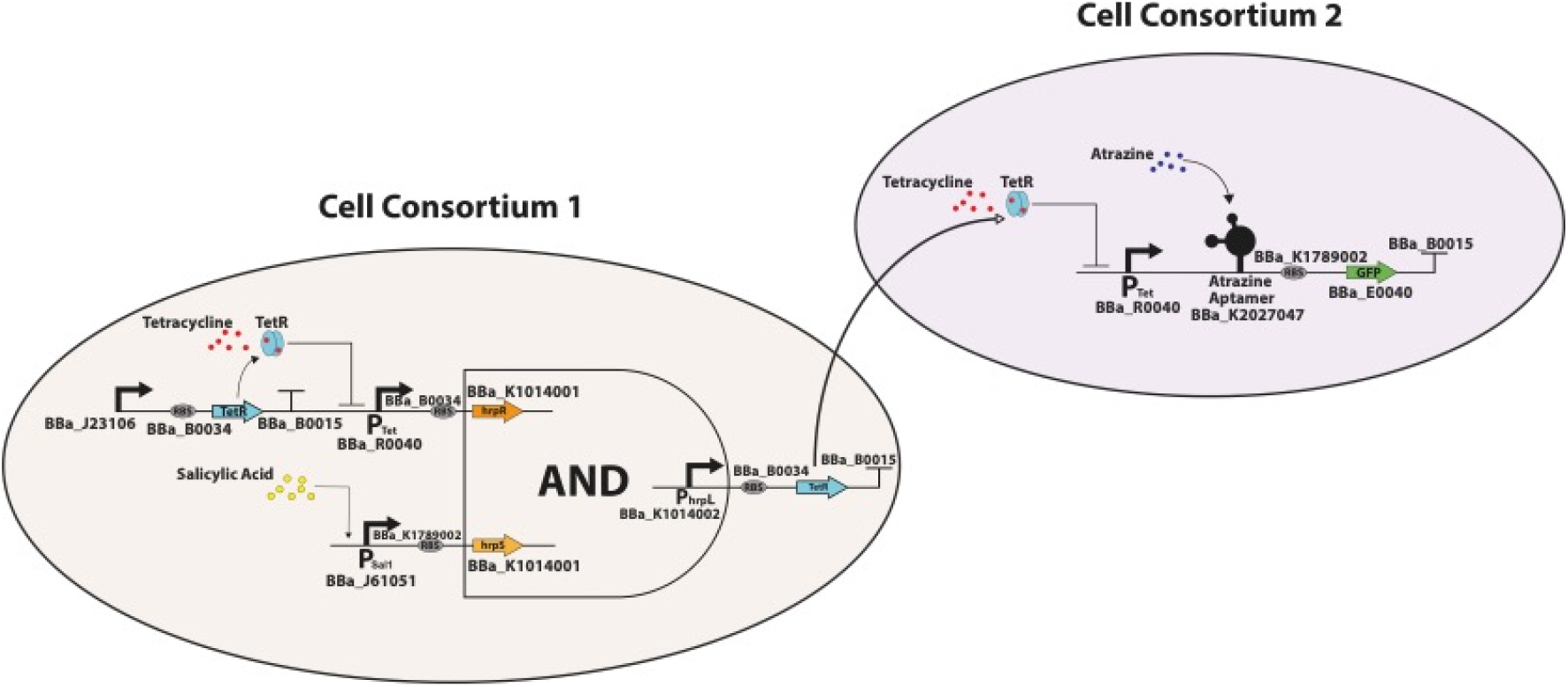
Schematic of the genetic circuits for the triple-input-AND-gated biosensor model. Two specific input signals (tetracycline and salicylic acid) are detected by the cell consortium 1 releasing TetR as an output. The cell consortium 2 detects the TetR released from the cell consortium 1 whilst also detecting atrazine to produce a GFP output.

#### 3. Toggle switch biosensor

The genetic toggle switch biosensor is constructed using two repressors (TetR and Salicylic acid repressor), two constitutive promoters (P_Tet_ and P_Sal1_), and two reporter genes (GFP and RFP) (Figure 5). The promoters are able to inhibit each other through the repressor proteins they code for. This design was selected as it requires the fewest genes and cis-regulatory elements theoretically to achieve a robust bistable behaviour [50]. The bistability of a genetic toggle switch arises from the mutually inhibitory set up of the repressor genes. Although the previous two biosensor designs were able to detect all three CECs simultaneously, the genetic toggle switch can detect only two contaminants at a time. The toggle switch biosensor is, thus, modelled to detect tetracycline (Figure 5A) and salicylic acid (Figure 5B). Fig. 5A illustrates state 1 of the genetic toggle switch, when a low amount of RFP will be produced by the activation of P_Sal1_ promoter. This state is altered in the presence of tetracycline molecules which binds to TetR. Consequently, the affinity between TetR and P_Tet_ reduces, resulting in the production of salicylic acid repressor proteins and GFP. Subsequently, the salicylic acid repressor proteins repress the P_Sal1_ promoter, preventing further production of TetR proteins. This results in a change in the state of toggle switch to state 2 as shown in Fig. 5B. In order to go back to state 1, salicylic acid must be detected by the sensor to prevent the repression of P_Sal1_; thus, increasing the production of TetR and RFP. Production of red fluorescent signals, therefore, indicates salicylic acid detection, while tetracycline presence is indicated by green fluorescent signals.

**Figure 5.**
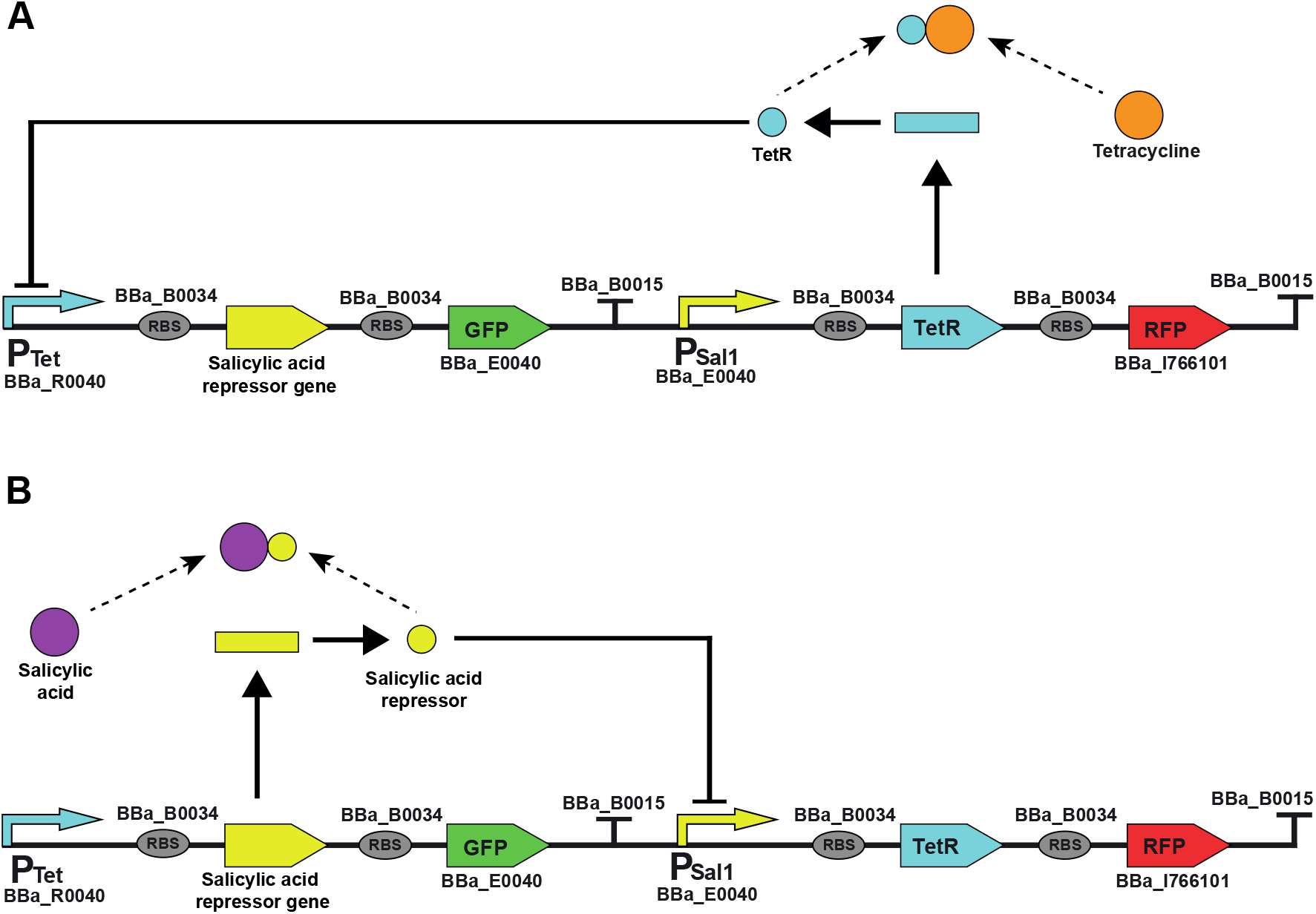
Schematic of the genetic circuits for the toggle switch biosensor model. (A) shows state one of the genetic toggle switch, in which TetR and RFP are produced. TetR represses P_Tet_ promoter preventing production of salicylic acid repressor gene and GFP. (B) shows state two of the genetic toggle switch, in which salicylic acid repressors and GFP are produced. Salicylic acid repressor represses the P_Sal1_ promoter preventing further production of TetR and RFP.

In this model, the protein degradation rates for GFP and RFP are assumed to be much higher than the degradation rates of TetR and salicylic acid repressor proteins. This assumption was made to ensure that the retention time of the TetR and salicylic acid repressor molecules would be much higher in comparison to GFP and RFP. Therefore, sufficient amounts of relevant repressor proteins would be ever-present in the system to bind with the inducers. More research is needed to identify the salicylic acid and atrazine repressor proteins to develop the correct parts to incorporate salicylic acid and atrazine detection in the toggle switch.

### Simulation and Analysis of Biosensor Models

#### 1. Ternion biosensor

The ODE simulation results (Figure 6) of the ternion biosensor model are shown in Fig. 6A. Initially, the biosensor was at ‘off’ state as no contaminant was present in the system. Tetracycline was introduced to the system at 1,000 seconds. This gave rise to a sharp increase in the concentration of YFP which plateaued at a value of 60 molecules. Once tetracycline was removed from the system, the concentration of YFP decreased to 0 molecules, showing the dependency of YFP production on tetracycline levels. At 1,100 seconds a constant influx of atrazine was introduced into the system and was maintained for a period of 1,000 seconds. This resulted in a sharp rise in the concentration of GFP which eventually plateaued at 1,750 seconds with a concentration of 250 molecules. Once atrazine was removed from the system, the GFP protein levels began to diminish over time. This illustrates the proportionality of the GFP output to atrazine levels. The introduction of salicylic acid into the system at 1,200 seconds gave a similar response to that of atrazine. Once salicylic acid was introduced into the system, RFP was rapidly produced and started to level out and reach steady state around 1,875 seconds also at a concentration of 250 molecules. Once salicylic acid was removed, the RFP proteins began to diminish over time due to its degrading nature. This sensor response illustrates the proportionality of the RFP output to the salicylic acid levels. The simulation results for the atrazine and salicylic acid sensors showed an equal amount of their respective reporter proteins being produced, however, the tetracycline sensor model produced much less of its reporter protein (Fig. 6A). This discrepancy may be due to the amount of TetR present in the system, brought about by the constitutive promoter BBa_J23106, repressing P_Tet_. This is an indication of the different output responses which may occur due to the mechanisms by which the reporter proteins are produced.

**Figure 6.**
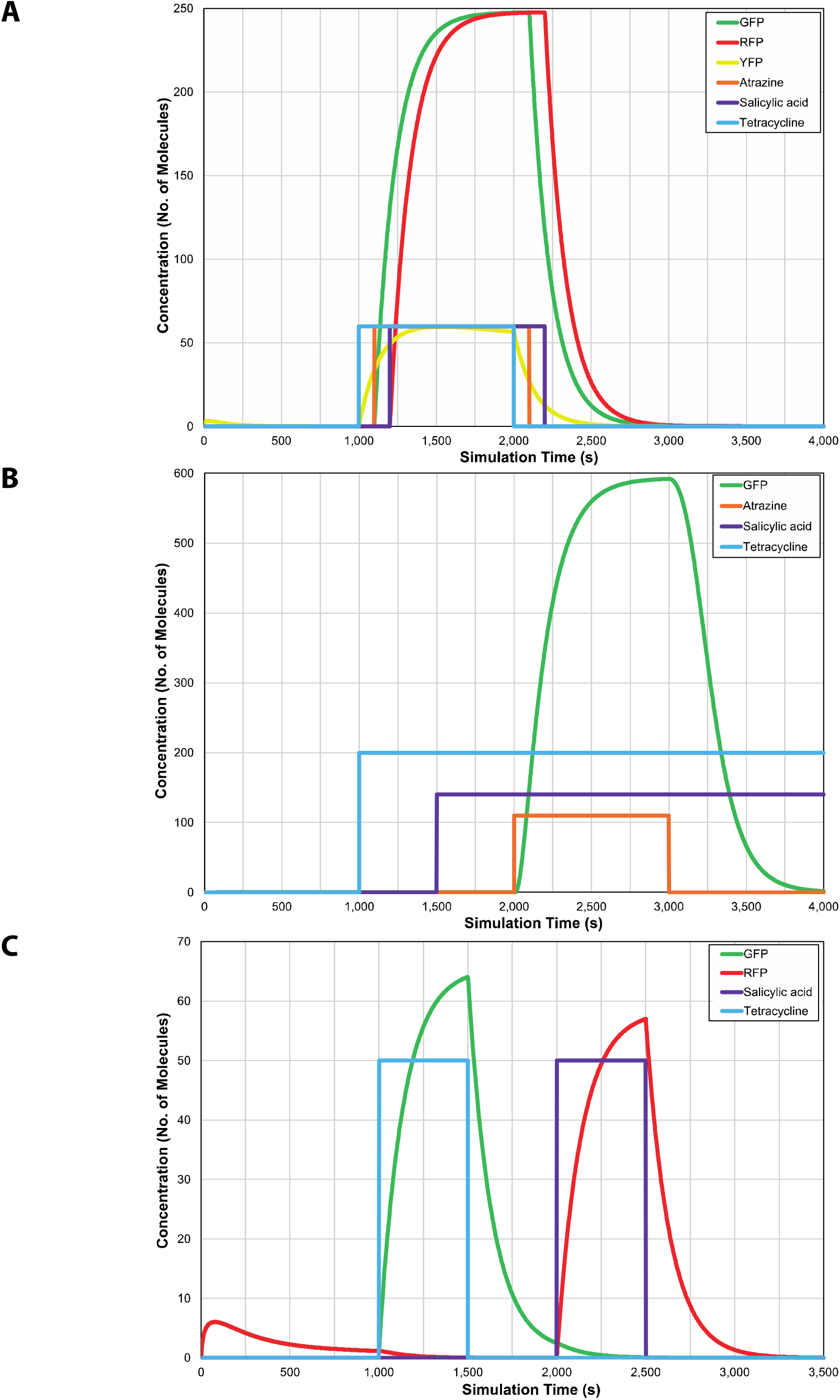
ODE simulation of biosensor models. (A) ODE simulation results of the ternion biosensor, (B) ODE simulation results of the triple-input-AND-gated biosensor, and (C) ODE simulation results for the toggle switch biosensor.

#### 2. Triple-input-AND-gated biosensor

Fig. 6B shows the simulation results for the triple-input-AND-gated biosensor model. As the biosensor produces only a single green fluorescent output from the detection of three CECs, no GFP production initially indicated at its ‘off’ state. Despite the introduction of tetracycline and salicylic acid at 1,000 seconds and 1,500 seconds, respectively in the system, GFP was not produced due to the AND-logic of the circuit was not satisfied. However, the addition of atrazine at 2,000 seconds, along with the presence of tetracycline and salicylic acid into the system allowed the transcription process to begin for the production of GFP. Thus, the presence of all three CECs resulted in a rapid production of GFP, which reached a maximum concentration of 600 molecules before decreasing steadily. At 3,000 seconds, atrazine was removed from the system, which resulted in an instant reduction of GFP production and the green fluorescent signals. This response behaviour supports the logic character of the biosensor, as GFP is only produced in the presence of all three contaminants in the system.

#### 3. Toggle switch biosensor

The genetic toggle switch biosensor behaviour in response to CECs is shown in Fig. 6C. There was an initial rise of the concentration of GFP and RFP to 6 molecules before decreasing back to 0 molecules at 1,000 seconds was observed. This initial rise of reporter proteins occurred as a result of setting the promoter counts of P_Tet_ and P_Sal1_ to be greater than zero to ensure that the toggle switch was capable of detecting either tetracycline or salicylic acid; thus, resembling a NOR-logic gate. Fig. 6C also shows a rapid increase of GFP, reaching a maximum of 67 molecules, when 50 molecules of tetracycline were added to the system at 1,000 seconds. This is caused by the complex formation reaction that takes place between the TetR proteins and tetracycline molecules. Once tetracycline leaves the system at 1,500 seconds, the GFP output rapidly decreases. Following this, salicylic acid was introduced into the system at 2,000 seconds. Once salicylic acid is in the system, it begins to bind with the salicylic acid repressor proteins available, which consequently permits the transcription for RFP, producing a maximum molecular count of 57 RFP molecules. When salicylic acid is removed from the system at 2,500 seconds, the RFP diminishes over time. Although equal quantities of tetracycline and salicylic acid molecules were added, more GFP molecules were produced as compared to RFP molecules. This difference is due to the repression of P_Sal1_ by the traces of salicylic acid repressors already present in the system when salicylic acid was added.

The simulations of the biosensor models were accomplished using ODE analysis. This type of analysis makes two significant assumptions. One being that the reactions that are occurring discretely, are assumed to occur continually instead [50]. The other being that the concentration of molecules present in the system varies [50]. However, these assumptions vary greatly to what actually occurs in genetic circuits [50]. For this reason, the simulation results of the biosensor models may not fully represent the results that would occur in experimental testing and implementation of the biosensors. Thus, a better representation of experimental results would be possible through the implementation of stochastic chemical kinetics in biosensors models.

### Factors to consider for experimental testing and field use of biosensors

Despite all the simulation results indicating the viability of the various biosensor designs, further experimental testing is needed to increase their reliability. In order to conduct experimental testing on these biosensors, several factors need to be considered. In the whole-cell biosensors, it is recommended that the cells used are of the prokaryotic nature and are derived from the bacteria, *Escherichia coli*. *E. coli* variants have been extensively used in previous experimental implementation of biosensors [51]–[53]. Temperature and pH levels are also important factors that need to be considered. The simulated biosensor models did not consider optimum operating conditions for the biosensors to detect various contaminants. Although whole-cell biosensors are able to operate over a broad range of temperatures and pH levels, the optimum conditions can only be obtained through their experimental testing [54]. In conjunction with temperature and pH levels, horizontal gene transfer (HGT) is also an important factor which needs to be considered during experimental testing of the designed biosensors. HGT is interpreted as the movement of genetic material across organisms and has been shown to occur in *E. coli* strains [55], [56]. Consequently, the biosensors may be susceptible to HGT; thereby, affecting their competence and test results. HGT may also lead to serious environmental and health issues.

Another important factor which needs to be considered is the undesired crosstalk between cells. Biological crosstalk refers to an instance where one or more components of one signal transduction pathway affect another [57]. Crosstalk may occur through the overexpression of particular proteins [58]. Additionally, crosstalk may occur through the overexpression of an unnatural response regulator [58]. Ninfa *et al* [59] observed the overexpression of a mutant version of nitrogen regulator II (NRII) in *E. coli*. This resulted in the phosphorylation of CheY, a non-cognate phosphorylation receptor and undesired interactions through which the swimming behaviour of *E. coli* was affected [59]. Effects of crosstalk may be minimised through the use of automation [60]. Additional challenges such as the time required to generate signals by reporter genes may arise when employing logic gated biosensors in the field. When compared to the ternion biosensor, the triple-input-AND-gated biosensor has more genetic circuit layers, thus, may require more time to transcribe the reporter gene further downstream. Moreover, the diffusion of TetR across the cell consortium may delay the production of the reporter proteins. Due to the structural intricacy of two-component systems, these sensors could also fail for reasons beyond present knowledge [58]. Furthermore, the stability of logic gated biosensors may give rise to problems in the field use. Thus, experimental testing is required to examine the stability of this type of biosensors. The mechanism for the regulation of P_hrpL_ promoter by hrpS and hrpR is not yet fully understood [48]. Hence, research in this area will permit the development of techniques to increase the efficiency of promoter activation.

Although iBioSim allowed us to model and simulate different biosensor designs, it lacks the ability to predict their experimental responses and behaviours. Also, the stability of the genetic circuits in the biosensor model is currently unexplainable through iBioSim. Furthermore, the promoter strength and specificity of genetic circuits are not considered in the iBioSim model simulations. Thus, to better indicate experimental results, software with the ability to integrate models and the phenotypic behaviour of the circuits must be developed [50].

## Conclusion

Recent advances in synthetic biology have increased the potential for groundwater contamination detection through the use of whole-cell biosensors. One such advancement is the development of genetic circuits using standard biological parts that can perform various logic operations and regulatory functions using biological cells. This advancement is further aided by the development of computational synthetic biology tools. The availability of GDA software such as iBioSim now allows us to develop and analyse various design models of genetic circuits without any experimental implementation in a living system. Using iBioSim, three whole-cell biosensor models were designed and analysed to detect multiple groundwater CECs simultaneously. The viability of these biosensors has also been analysed and presented using mathematical models and simulation results. Despite these promising results, current use of whole-cell biosensors in environmental monitoring and field applications is significantly low. This is partly due to the lack of predictability and optimum operating conditions of whole-cell biosensors in field settings. Research gaps in transcriptional regulation of biosensor parts also need to be addressed to increase the predictability of these biosensors. Furthermore, computational synthetic biologists need to develop tools, which can imitate real life environments for more relevant simulations of biosensor models. With this framework in place, more reliable and robust whole-cell biosensor designs can be achieved for the detection of CECs in the future.

