## Supplemental file for "In Silico Design and Analysis of Genetic Circuit-Based Whole-Cell Biosensors for Detecting Groundwater Contaminants"

**iBioSim Sub Models**

**Ternion Biosensor**

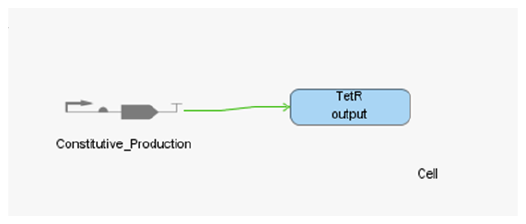

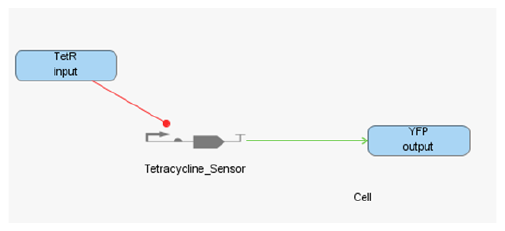

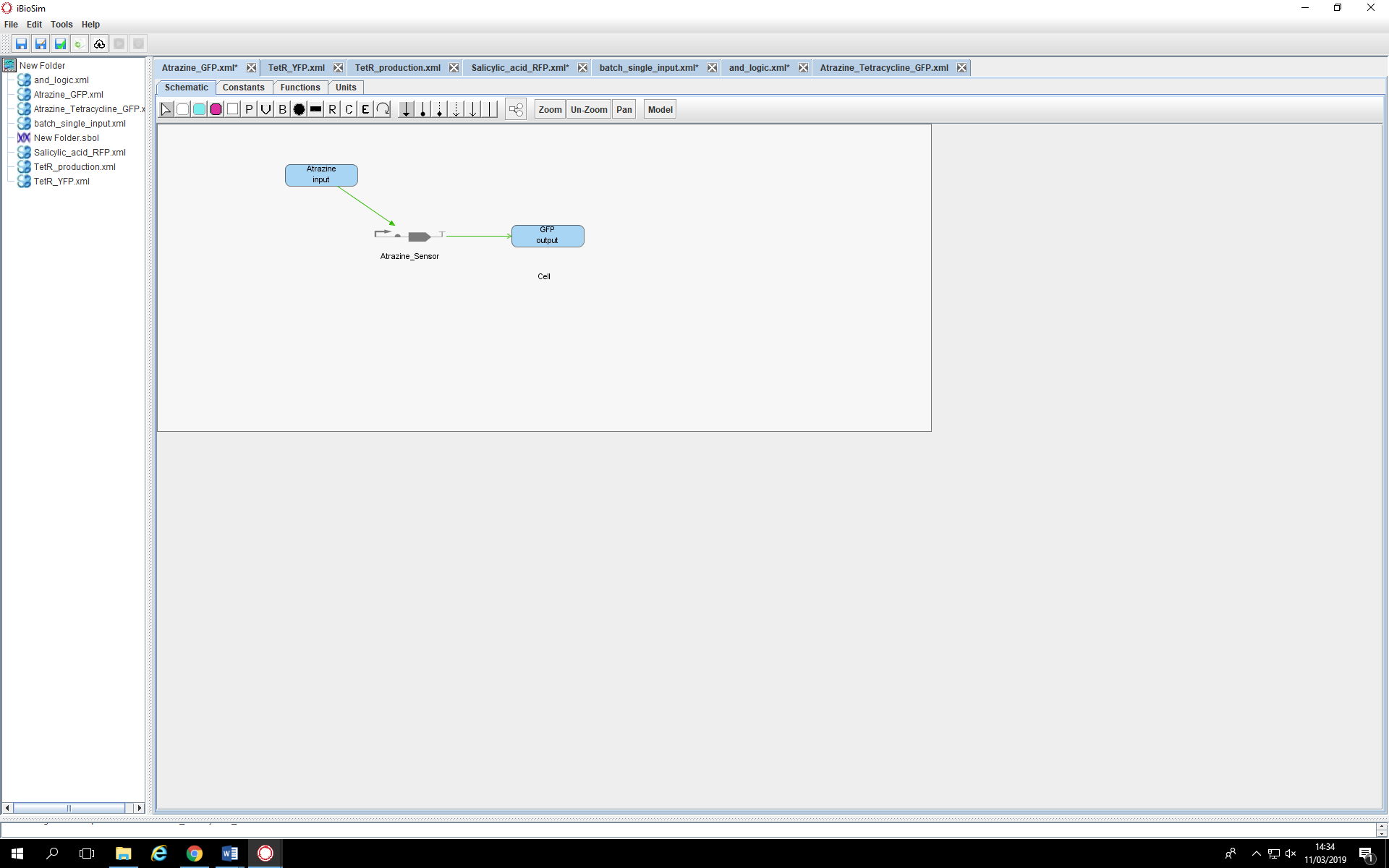

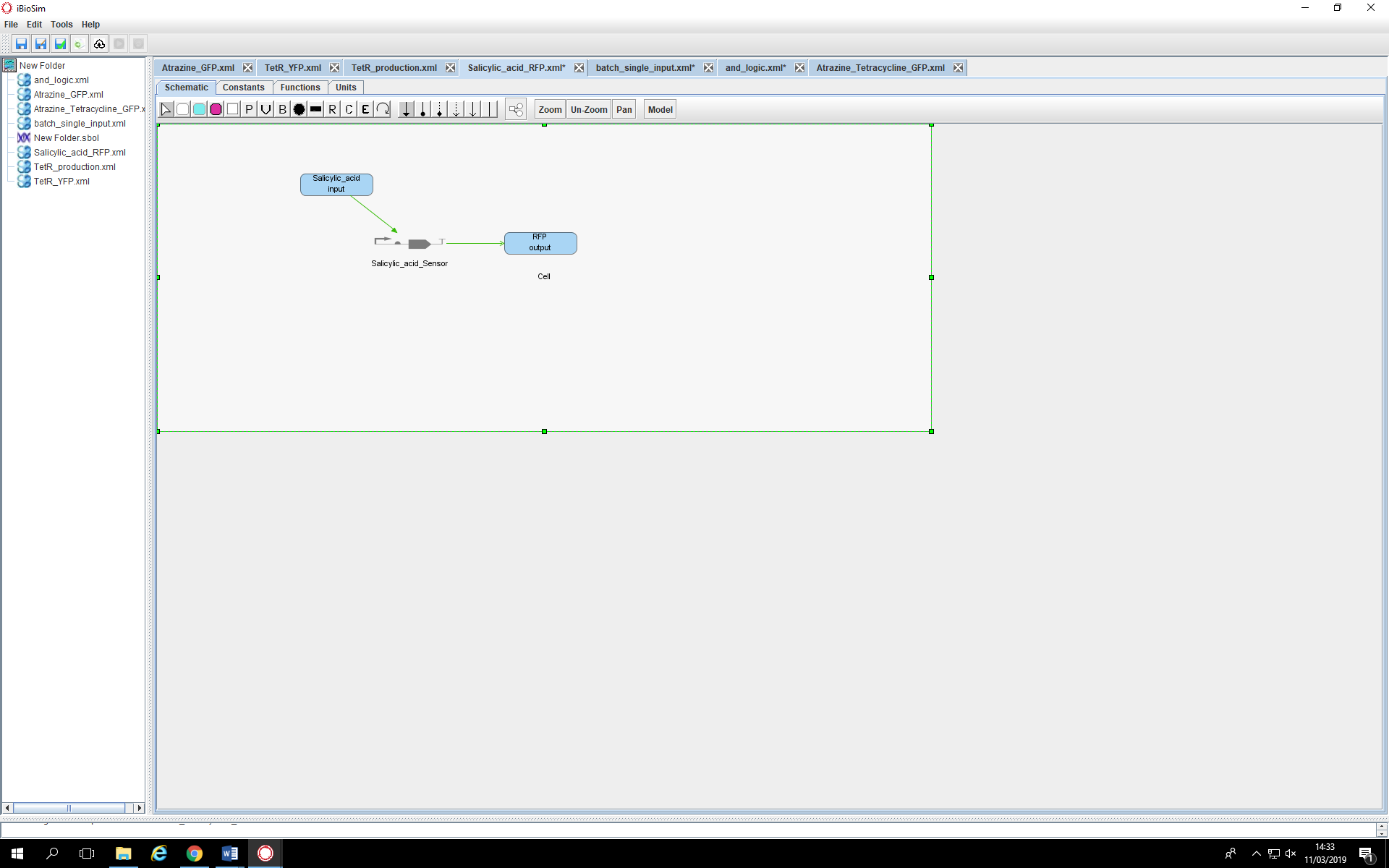

**Triple-Input-AND-Gated Biosensor**

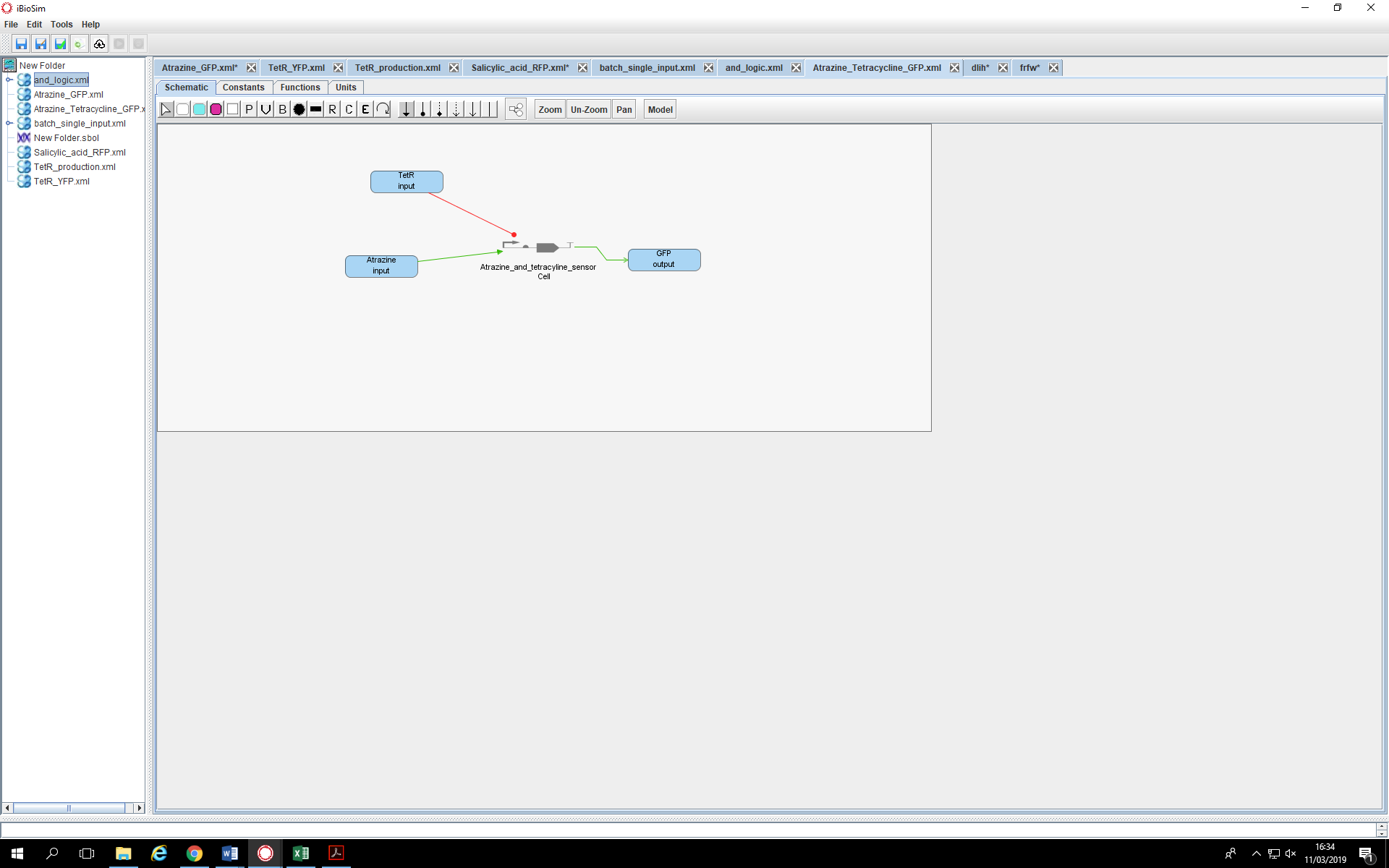

**Toggle Switch Biosensor**

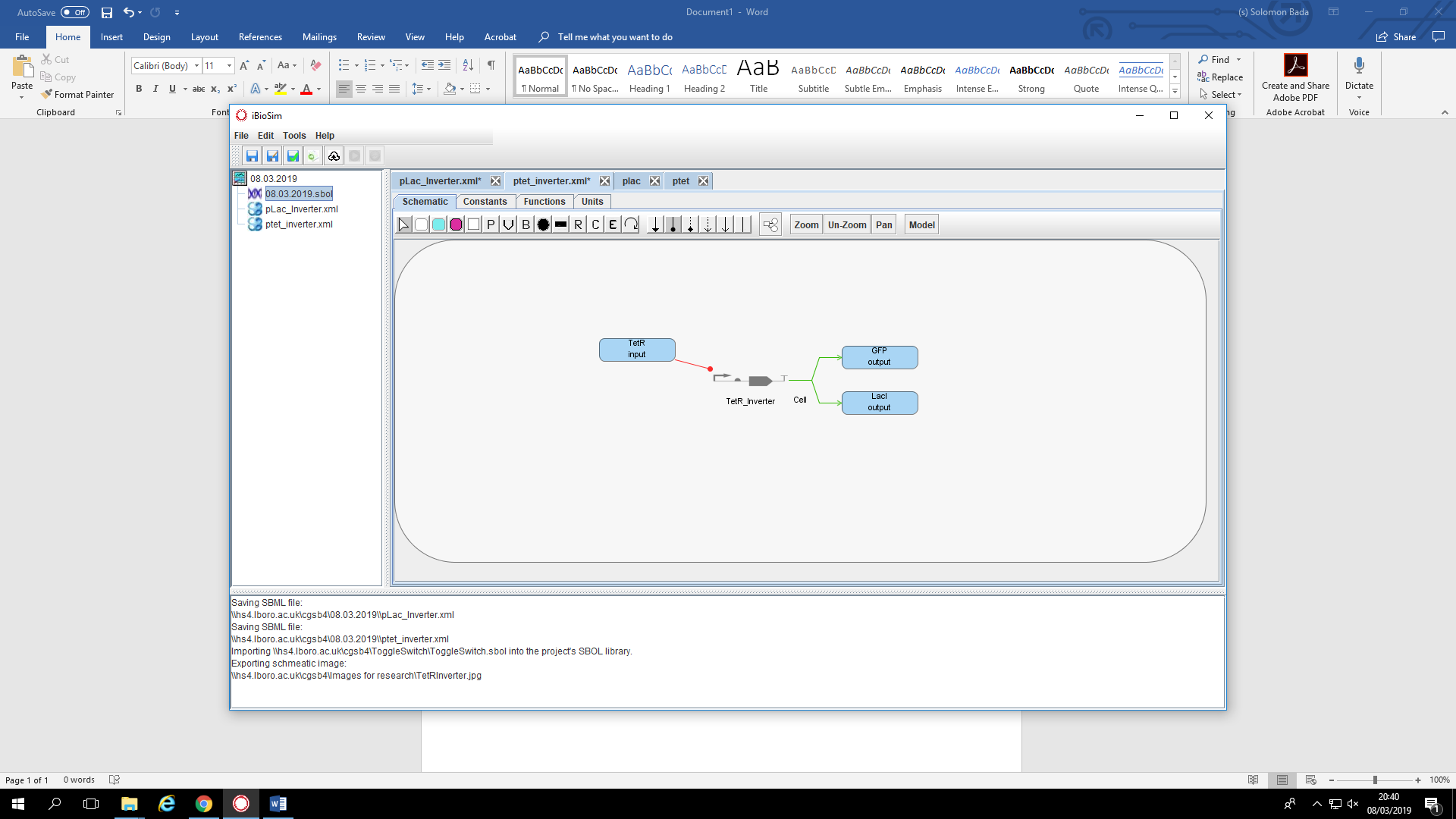

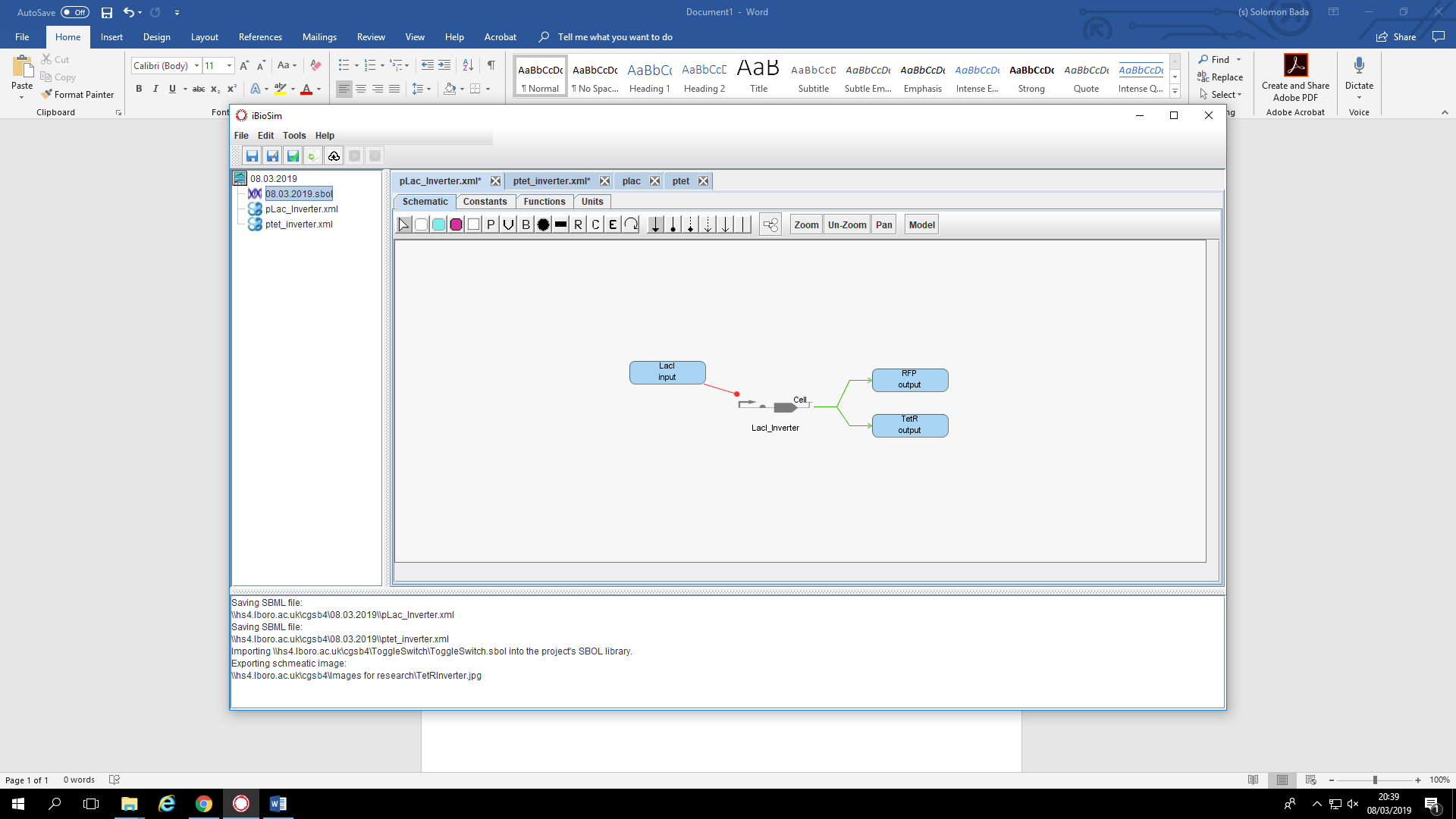

**SBOL Visual Design of Genetic Circuits Used in Different Biosensors**

**Ternion Biosensor**

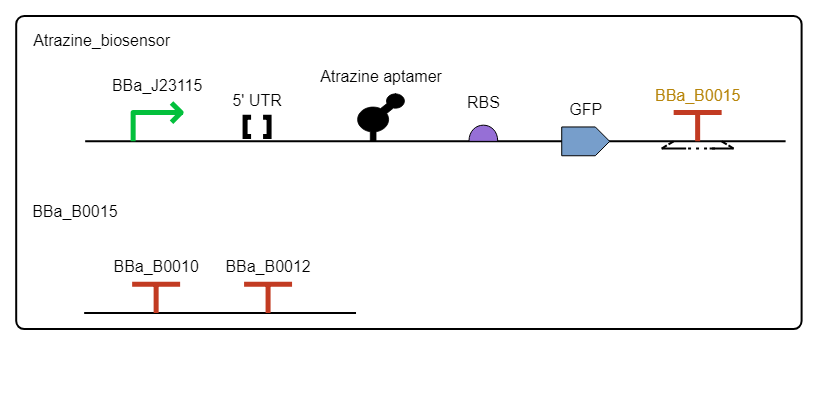

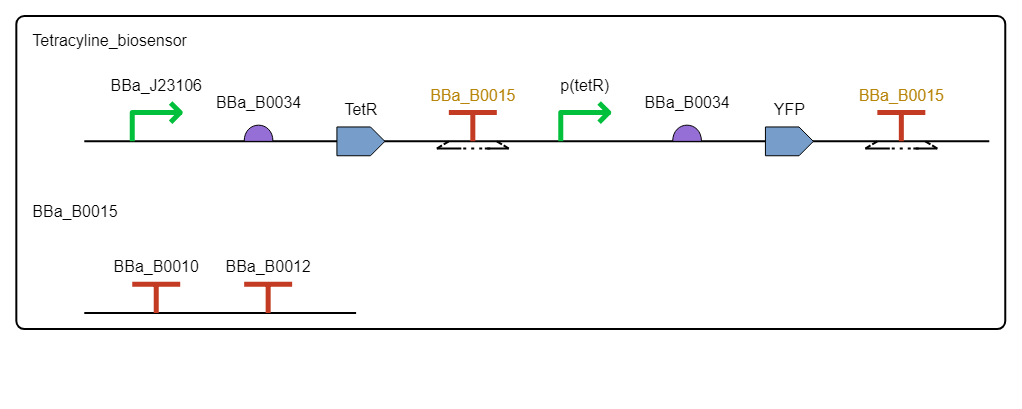

**Triple-Input-AND-Gated Biosensor**

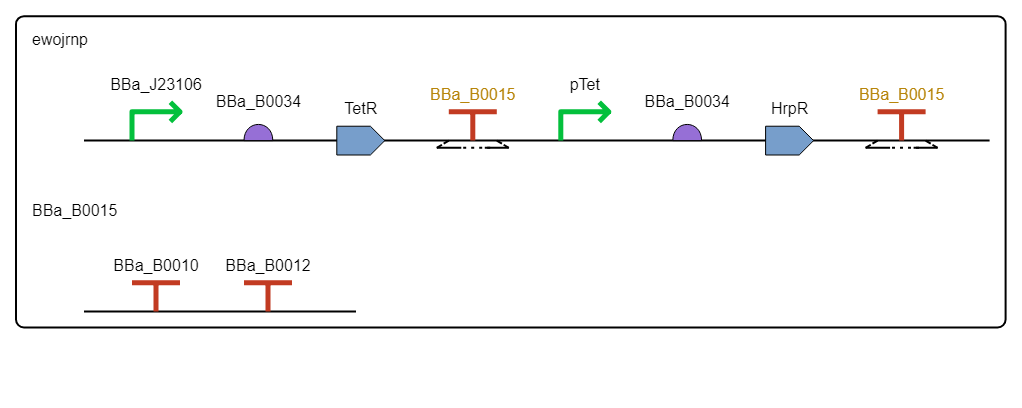

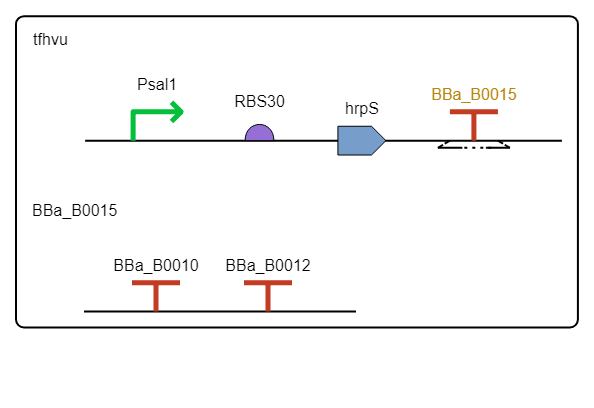

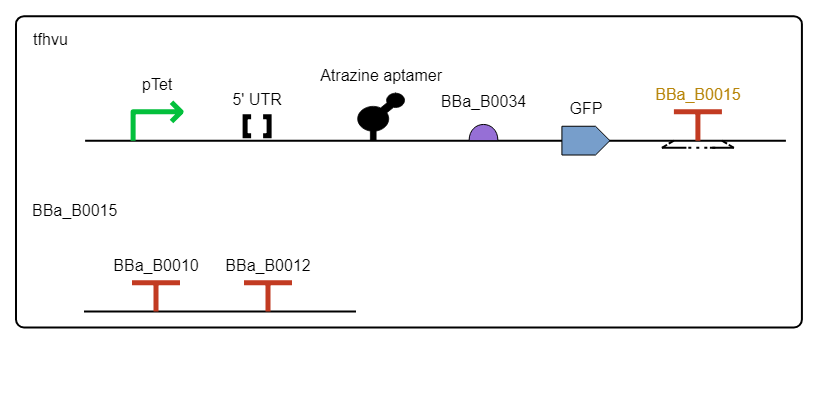

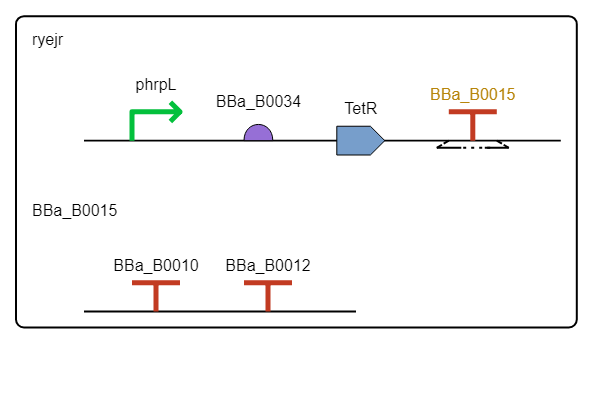

**Toggle Switch Biosensor**

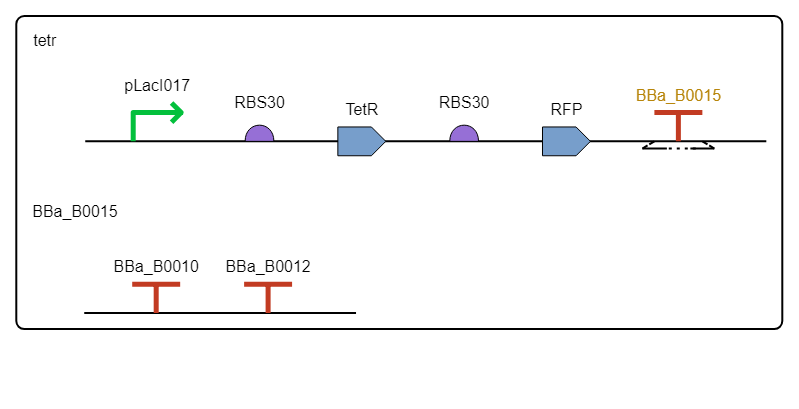

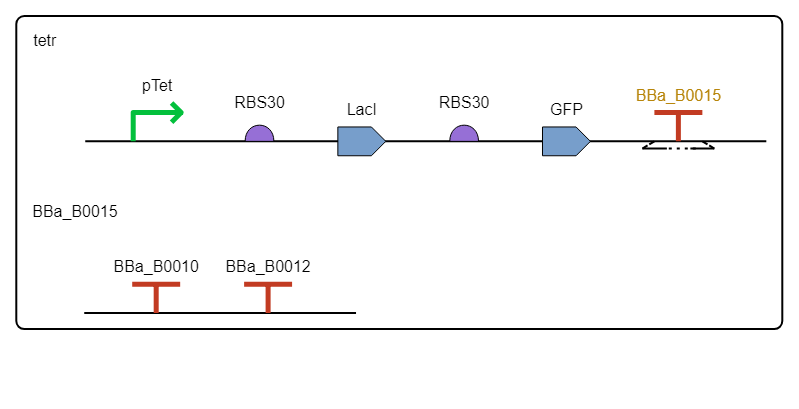

**Mathematical Derivation of Biosensor Model Equations**

**Ternion Biosensor**

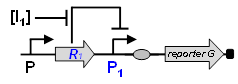

In the diagram, the expression of the protein 𝐺 is dependent on the presence of the inducer 𝐼1. The gene expression can be modelled as [1], [2]:

| $\frac{d[G]}{dt}\text{ = }\alpha\cdot k_{1}+\frac{k_{1}\cdot[I_{1}]^{n}}{[I_{1}]^{n}+{K_{1}}^{n}}-d\cdot[G]$ | A1 |
| --- | --- |

Where the concentration of the inducer is show as $I_{1}$.The promoter’s basal activity is shown as $\alpha\cdot k_{1}$ The protein degradation activity is shown as $d\cdot[G]$. The Hill constant $K_{1}$ and coefficient $n$ account for the interaction between the promoter-regulator and inducer. $k$ is the maximum rate of expression as a result of induction.

At steady state Eq. A1 becomes:

| $f\left( \left[ I_{1} \right] \right)=\left[ G \right]_{ss}=k(\frac{\alpha+\left[ I_{1} \right]^{n}}{K_{1}^{n}+\left[ I_{1} \right]^{n}})$ | A2 |
| --- | --- |

The steady state degradation activity of protein G is shown as $\left[ G \right]_{ss}$.

**Triple-Input-AND-Gated Biosensor**

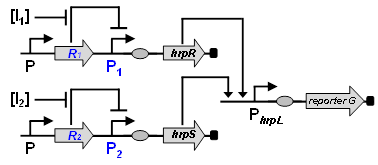

The diagram illustrates the mechanism of an AND gated biosensor. Expression of the protein G is regulated by the hrpL promoter P_hrpL_. P_hrpL_ is activated by the presence of hrpL which is formed by the hetero proteins hrpR and hrpS. These proteins are made in the presence of the inducers $I_{1}$ and $I_{2}$. Therefore, the transfer function can be shown as the product of the two hill functions curves [1], [2]. The gene expression for the triple-input-AND-gated biosensor can be modelled by:

| $f\left( \left[ R \right]_{ss}, \left[ S \right]_{ss} \right)=\frac{\left[ G \right]_{ss}}{\left[ G \right]_{ssmax}}=\frac{{(\left[ R \right]_{ss}/K_{R})}^{n_{R}}{(\left[ S \right]_{ss}/K_{S})}^{n_{S}}}{(1+{(\left[ R \right]_{ss}/K_{R})}^{n_{R}})(1+{(\left[ S \right]_{ss}/K_{S})}^{n_{S}})}$ | A3 |
| --- | --- |

The letters $R$ and $S$ correspond to the hrpR and hrpS proteins, respectively. The AND gate maximum output level at steady state is given as $\left[ G \right]_{ssmax}$. The Hill constant $K$ and coefficient $n$ account for the interaction between the promoter-regulator and inducer.

**Genetic Toggle Switch Biosensor**

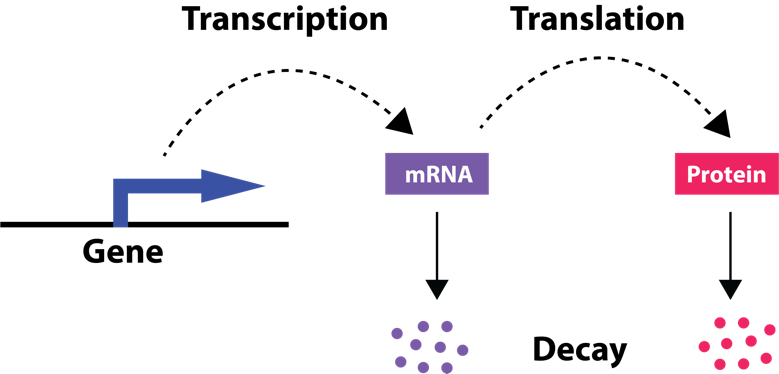

The scheme above can be translated into mathematical equations, which can be seen below.

| $\frac{d[mRNA]}{dt}=k_{transcription}\left[ DNA \right]-\gamma_{m}[mRNA]$ | A4 |
| --- | --- |

| $\frac{d[G]}{dt}=k_{translation}\left[ mRNA \right]-\gamma_{p}$ | A5 |
| --- | --- |

For a toggle switch, before the equation can be made, the toggle switch must be reduced into a one expression system independently.

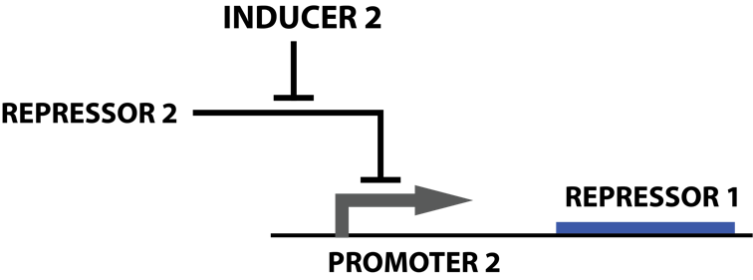

The production of mRNA relies on the transcription constant and activator or repressor concentration.

| $Chemical reaction:\left[ R \right]+\left[ P \right]\begin{matrix} k_{on} \\ \leftrightarrow\\ k_{off} \end{matrix}[RP]$ | A6 |
| --- | --- |

| $Bound fraction=\left( \frac{K[R]}{1+K[R]} \right)^{n}$ | A7 |
| --- | --- |

In the repression model, the following is incorporated:

| $Unbound fraction=1-bound fraction= \left( \frac{1}{1+K[R]} \right)^{n}$ | A8 |
| --- | --- |
| $Promoter activity= K_{transcription}* Unbound fraction=\frac{K_{transcription}}{1+\left( K[R] \right)^{n}}$ | A9 |

In the toggle switch, the promoter activity depends on the inducer concentration also, due to the inducer’s ability to inhibit repressor to repress a promoter.

| $\left[ I \right]+\left[ R \right]\begin{matrix} k_{on'} \\ \leftrightarrow\\ k_{off'} \end{matrix}[IP]$ | A10 |
| --- | --- |
| $Unbound fraction= \left( \frac{1}{1+K^{'}[I]} \right)^{n'}=\frac{[R]}{\left[ IR \right]+[R]}$ | A11 |
| $\left[ R \right]=\frac{[IR]}{\left( K^{'}[I] \right)^{n'}}$ | A12 |
| $\frac{d\left[ mRNA \right]}{dt}=k_{transcription}\left[ DNA \right]-\gamma_{m}\left[ mRNA \right]$ | A13 |
| $Promoter Acitivity= \frac{K_{transcription}}{1+\left( K[R] \right)^{n}}$ | A14 |
| $\left[ R \right]=\frac{[IR]}{\left( K^{'}[I] \right)^{n'}}$ | A15 |
| $\frac{d\left[ mRNA \right]}{dt}=\frac{K_{transcription}}{1+K\left( \frac{\left[ IR \right]}{\left( K^{'}\left[ I \right] \right)^{n^{'}}} \right)^{n}}-\gamma_{m}[mRNA]$ | A16 |

For protein production rate, the transcription rate must be multiplied with the number of molecules of protein product.

| $protein production rate=P*\frac{K_{transcription}}{1+K\left( \frac{[IR]}{\left( K^{'}[I] \right)^{n'}} \right)^{n}}$ | A17 |
| --- | --- |

The degradation time of protein depends on concentration of protein and its lifetime.

| $protein degradtion rate=[G]$ | A18 |
| --- | --- |

From the following equations above, a differential equation of protein product as a function of time can be formulated.

| $\frac{d[G]}{dt}=P*\frac{K_{transcription}}{1+K\left( \frac{\left[ IR \right]}{\left( K^{'}\left[ I \right] \right)^{n}} \right)^{n^{'}}}-[G]$ | A19 |
| --- | --- |

| **Parameter** | **Description** |
| --- | --- |
| [DNA] | Concentration of DNA |
| $\left[ I \right]$ | Free Inducer |
| $\left[ IR \right]$ | Inducer-Repressor Complex |
| [mRNA] | Concentration of mRNA |
| $n$ | Hill coefficient |
| [G] | Protein degradation rate |
| $\left[ P \right]$ | Free Promoter |
| $P$ | Number of molecules of protein product |
| $\gamma_{m}$ | mRNA degradation rate |
| $\gamma_{p}$ | Protein degradation rate |
| $\left[ R \right]$ | Free Repressor |
| $k_{transcription}$ | Rate of RNA synthesis |
| $k_{translation}$ | Rate of protein synthesis |
| $\left[ RP \right]$ | Repressor-Promoter Complex |
| $T$ | Protein lifetime |
| $K$ | $\frac{k_{on}}{k_{off}}$ |
